# Restored TDCA and Valine Levels Imitate the Effects of Bariatric Surgery

**DOI:** 10.1101/2021.01.11.425291

**Authors:** Markus Quante, Jasper Iske, Timm Heinbokel, Bhavna N. Desai, Hector Rodriguez Cetina Biefer, Yeqi Nian, Felix Krenzien, Hirofumi Uehara, Ryoichi Maenosono, Haruhito Azuma, Johann Pratschke, Christine Falk, Tammy Lo, Eric Sheu, Ali Tavakkoli, David Perkins, Maria-Luisa Alegre, Alexander S. Banks, Abdallah Elkhal, Stefan G. Tullius

## Abstract

Obesity is widespread and linked to various co-morbidities. Bariatric surgery has been identified as the only effective treatment, promoting sustained weight loss and the remission of co-morbidities.

We performed sleeve-gastrectomies (SGx) in a pre-clinical mouse model of diet-induced obesity (DIO), delineating the effects on long-term remission from obesity. SGx resulted in sustained weight loss and improved glucose tolerance. Mass-spectrometric metabolomic profiling revealed significantly reduced systemic levels of taurodeoxycholic acid (TDCA) and L-valine in DIO mice. Notably, TDCA and L-Valine levels were restored after SGx in both human and mice to levels comparable with lean controls.

Strikingly, combined systemic treatment with TDCA and valine induced a profound weight loss in DIO mice analogous to effects observed after SGx. Utilizing indirect calorimetry, we confirmed reduced food intake as causal for TDCA/valine-mediated weight loss via a central inhibition of the melanin-concentrating hormone.

In summary, we identified restored TDCA/valine levels as an underlying mechanism of SGx-derived effects on weight loss. Of translational relevance, TDCA and L-valine are presented as novel agents promoting weight loss while reversing obesity-associated metabolic disorders.

## Introduction

Obesity is a global epidemic with broad clinical and economic consequences. Data by the Global Burden of Disease Study (GBD) estimate that overweight and obesity are causing 3.4 million deaths and 3.8 % of disability-adjusted life-years (DALYs) worldwide(*1*). Moreover, obesity is promoting numerous disorders including diabetes, cardiovascular disease (CVD) and cancer(*2, 3*). Therapeutic approaches including diets or approved pharmacological interventions treating obesity and related co-morbidities have only had limited success(*4, 5*).

Bariatric surgery including sleeve gastrectomies (SGx) have been successful in achieving a sustained body weight loss(*6*). Moreover, these procedures also display an effective treatment of obesity associated type 2 diabetes (T2D)(*7*) with durable HbA1C remissions (*8*), improved insulin sensitivity and glycemic control(*9*), all resulting into long-term reduction of overall mortality and obesity related risk factors(*5, 10*).

SGx has been conceptualized to elicit weight loss by physically restricting gastric capacity(*11*). Moreover, the procedure leads to a weight-independent reduction in plasma triglyceride levels(*12*), improved HDL levels(*13*), increased hepatic insulin sensitivity(*14*) and weight-independent alterations of gut-derived anti-diabetic hormones (GLP-1 and PYY(*15*)), implicating an altered metabolic profile.

At a molecular level, bile acid signaling is thought to constitute a mechanistic underpinning of bariatric surgery-induced weight loss. Bile acids have been shown to activate nuclear transcription factors involved in hepatic glucose metabolism(*16*) that promote GLP-1 secretion via signaling by the bile acid receptor TGR5(*17*). Moreover, bile acids have been identified to be essential for SGx-derived weight loss through signaling via the bile acid receptor FXR(*18*). Consistently, various studies have reported increasing serum bile acids following SGx in experimental and clinical studies(*19, 20*).

Thus, durable weight loss and amelioration of obesity-associated disorders after SGx are based on a complex metabolic and hormonal homeostatic circuitry rather than surgery-induced malabsorption alone. Clearly, understanding the mechanisms of bariatric surgery in detail and achieving weight loss and metabolic changes through a non-surgical treatment may represent a desired alternative.

Here, we made use of DIO mice, a well-established model of obesity and performed SGx to delineate how bariatric surgery promotes long term metabolic changes and the reduction of body weight.

Through an unbiased quantitative metabolomic analysis, we identified restored levels of taurodeoxycholic acid (TDCA), and L-valine following SGx in our experimental model and confirmed similar effects in patients undergoing SGx.

Administration of both, TDCA and valine in DIO mice caused a robust and sustained weight loss, reduced fat tissue and reversed DIO-associated T2D. Notably, indirect calorimetry using a Comprehensive Laboratory Monitoring System (CLAMS) demonstrated a drastically decreased food intake in the absence of reduced energy expenditure.

Dissecting hypothalamic neuroendocrine peptides associated with the regulation of appetite pointed towards the critical role of the hypothalamic orexigenic peptide melanin-concentrating hormone (MCH). Indeed, fasting untreated DIO mice, exhibited dramatically increased MCH levels when compared to TDCA/valine-treated animals. Consistently, TDCA/valine treatment combined with an intrathecal injection of recombinant MCH reversed the effects of TDCA/valine on weight reduction, thus confirming the involvement of MCH inhibition as the mechanistic path of TDCA/valine-induced weight reduction.

## Results

### Bariatric surgery induces sustained weight loss by restoring the metabolite profile in both, mice and humans

Bariatric surgery is effective in 80-90% of obese individuals and leads to sustained weight loss and significant improvement of co-morbidities(*5, 6, 10*). To assess mechanisms of surgically induced weight loss through Sleeve Gastrectomies (SGx), we made use of C57BL/6 wild type diet-induced obese (DIO) mice, a well-established murine model of obesity(*21, 22*) (**Fig. 1A)**. Subsequent to SGx, we observed a significant weight loss (−34% by 2 weeks). In marked contrast, sham-operated DIO mice re-gained their preoperative weight by postoperative day 14 (−5%), indicating that the observed weight loss was independent of the surgical trauma (**Fig. 1B)**. Thus, SGx promoted durable weight loss independent of the surgical procedure or dietary effects. Previous studies implicated that SGx induce modifications in levels of metabolic and intestinal hormones(*23*). Therefore, we performed a quantitative metabolomic profiling to characterize the consequences of obesity and weight-loss surgery on endogenous metabolites. Our unbiased approach detected 260 metabolites with 17 candidates showing a significant statistical difference between lean and DIO mice by analysis of variance (ANOVA), heat mapping and significance analysis of microarrays (SAM) **(Fig. 2A, B)**.

**Figure 1.**
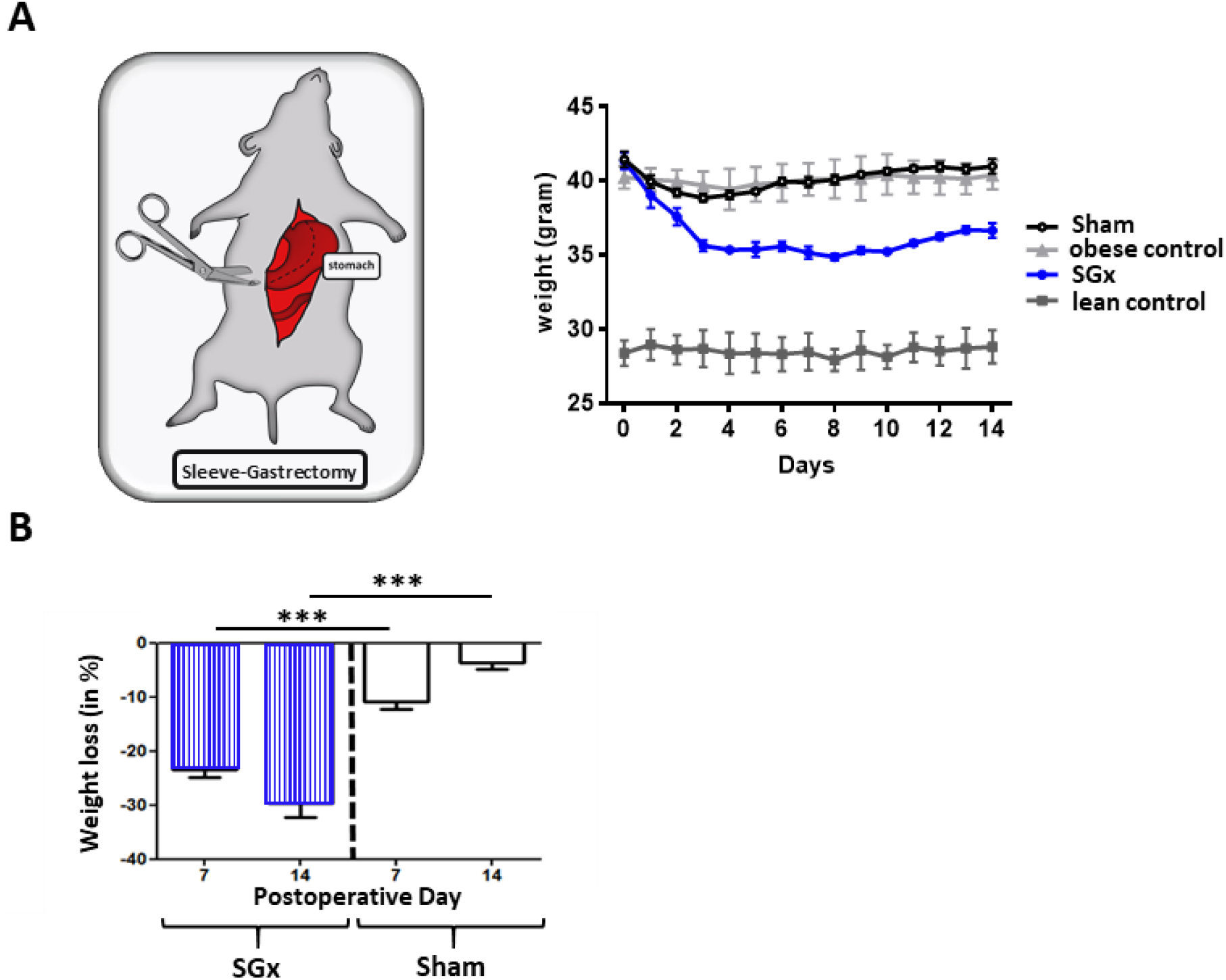
Sleeve gastrectomy induces significant weight loss independent of the surgical procedure. C57BL/6 DIO mice (n=5) underwent bariatric SGx while laparotomy was performed on animals of the sham group. Another set of C57BL/6 DIO and lean mice served as controls. **(A)** Body weight was monitored for a course of 2 weeks every 24 hours. **(B)** C57BL/6 DIO mice were returned to NCD in parallel to monitoring of SGx and Sham animals comparing mean weight loss after 7 and 14 days, respectively. Results are representative of at least three independent experiments. Column plots display mean with standard deviation. Statistical significance was determined using Two-Way-ANOVA followed by Turkey’s multiple comparison test with single pooled variance. Asterisks indicate p-values * = p<0.05, **= p<0.01 and *** = p<0.001. Only significant values are shown. (n=7 animals/group).

**Figure 2.**
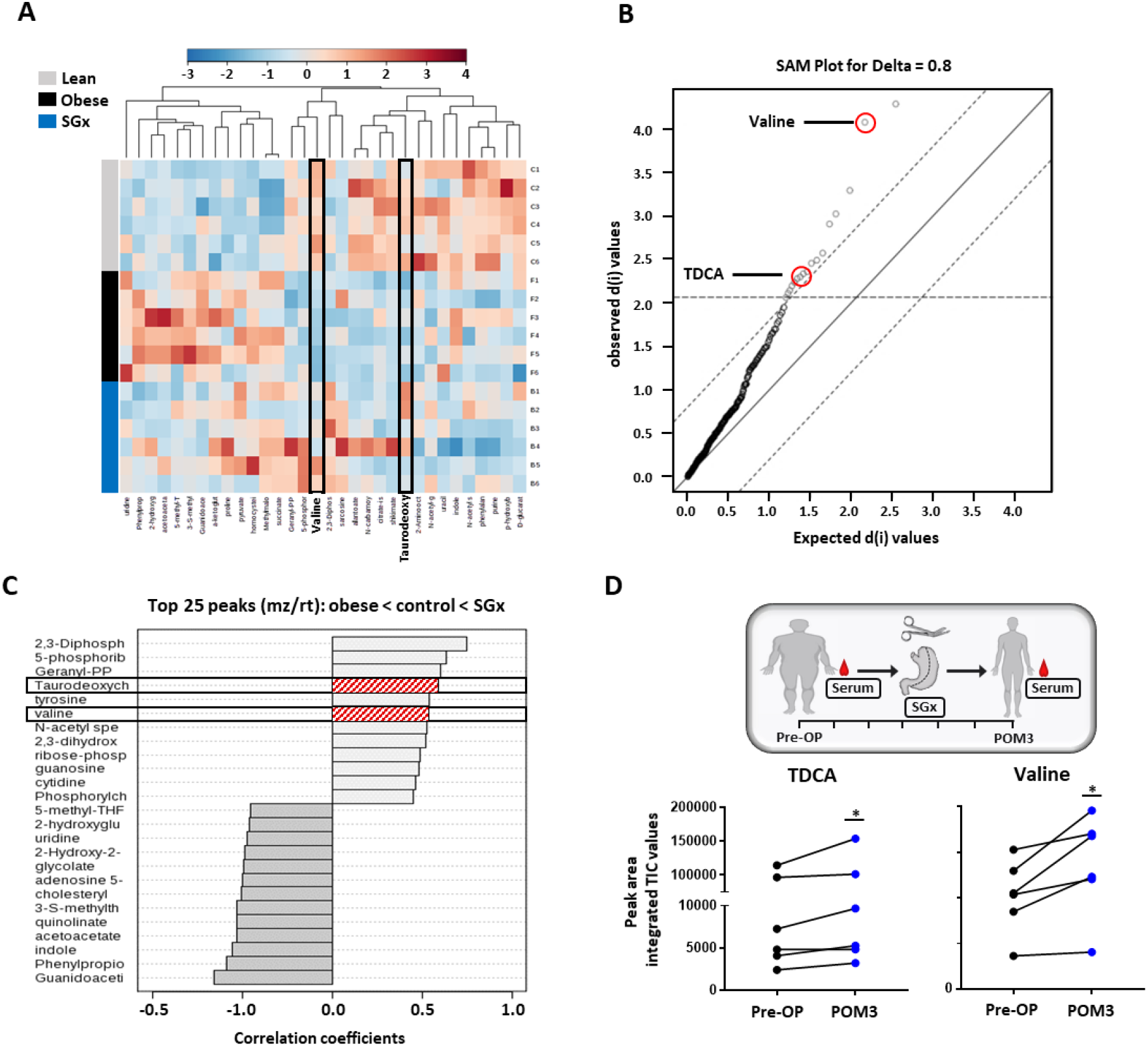
Sleeve gastrectomy restores systemic TDCA/valine levels in both DIO mice and obese humans. Whole blood samples from C57BL/6 DIO mice after SGx and DIO and lean controls were analyzed with a 5500 QTRAP mass spectrometer. Quantitative analysis was performed utilizing MetaboAnalyst 3.0. **(A)** Heatmap of 32 metabolites displayed after hierarchical clustering, p<0.05. **(B)** Significance analysis of microarrays (SAM) revealed 17 metabolites with significance **(C)** Pattern hunter stratified the 25 metabolites with top peaks (mz/rt) according to the order obese-control-SGx. **(D)** Serum was isolated from patients undergoing SGx pre-operative (Pre-OP) and 3 months after surgery (POM3). TDCA and valine levels were quantified using mass spectrometry and Peak are integrated TIC values compared (n=6). Results are representative of at least three independent experiments. Statistical significance was determined using One-Way-ANOVA and significance analysis of microarrays (SAM). TDCA/valine TIC values from human samples were compared using paired student’s T test. Asterisks indicate p-values * = p<0.05, ** = p<0.01, *** = p<0.001. Only significant values are shown. (n=6 animals/group, n=6 patients)

We found significantly decreased systemic levels of TDCA and L-valine in DIO mice. Obese animals, that underwent SGx, however, displayed overall restored levels of both metabolites, suggesting a critical involvement of TDCA/valine in the metabolic underpinning of SGx-induced weight loss. **(Fig. 2C)**.

Moreover, to investigate the translational relevance of our findings, we analyzed serum levels of TDCA and valine in human samples and collected from patients immediately prior to and three months after SGx **(Fig. 2D)**. Subsequent to clinical SGx, we observed a significant increase in both TDCA and valine levels, indicating a similar impact of SGx in mice and humans **(Fig. 2D)**.

### TDCA/Valine treatment induces robust weight loss and ameliorates obesity related insulin resistance

Based on our metabolomic profiling data, indicating restored systemic TDCA/valine levels after SGx, we next set out to assess the physiological impact of TDCA/valine on obesity and administered both metabolites intraperitoneally to naive DIO mice for a course of 2 weeks. The combined injection of TDCA and valine resulted in robust weight loss **(Fig. 3A)** that went beyond the observed effects in mice undergoing SGx **(Fig. 3B)**. Importantly, administration of TDCA/valine to lean control mice did not impact weight loss **(Fig. 3A)**.

**Figure 3.**
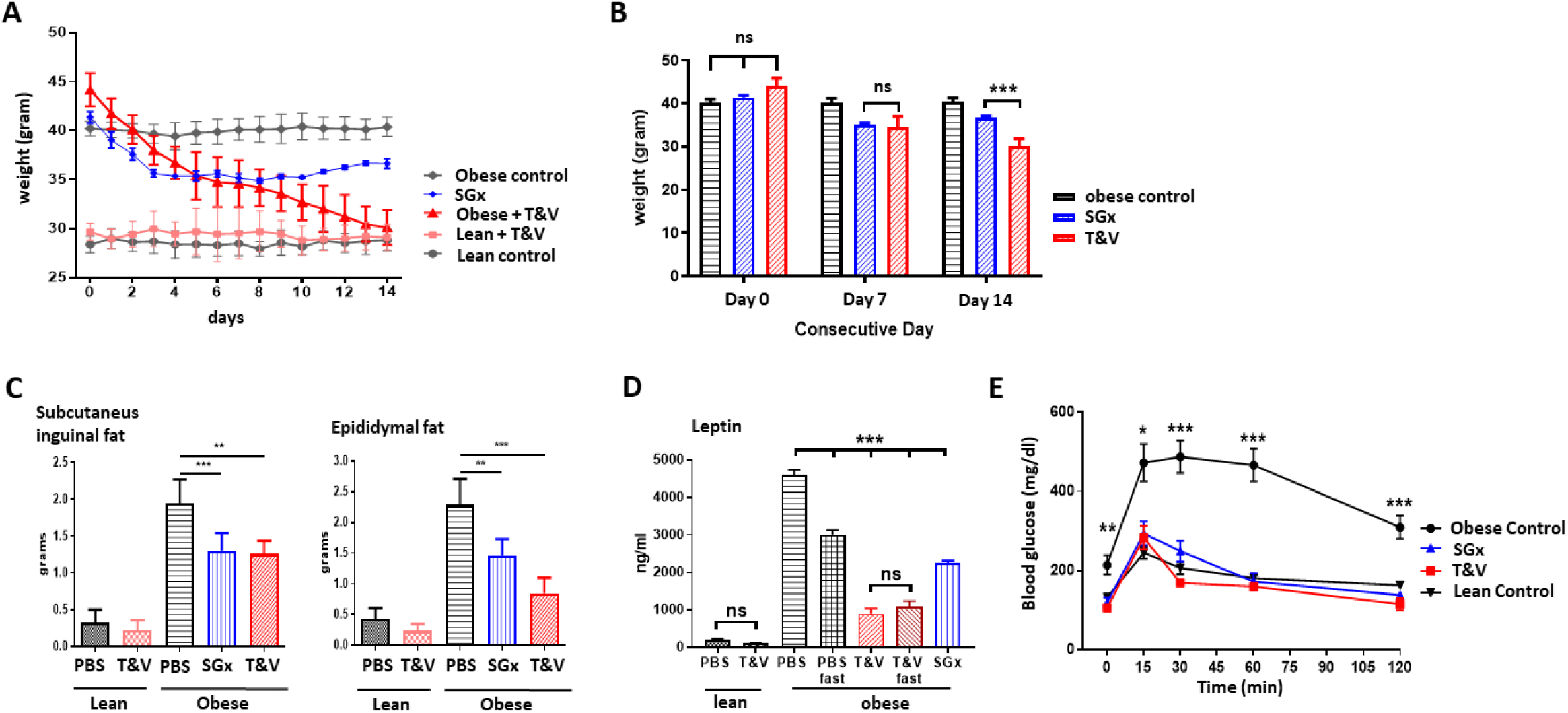
TDCA/Valine treatment induce robust weight loss and ameliorate obesity related insulin resistance. C57BL/6 DIO mice received intraperitoneal injections of TDCA (50mg/kg) and L-Valine (200mg/kg) daily over the course of 2 weeks. **(A)** Body weight was evaluated for 2 weeks every 24 hours. **(B)** Column plot of mean body weight comparing SGx and T/V-treated animals at day 0, 7, and 14. **(C)** Subcutaneous and epididymal fat tissue was removed after 14 days of treatment or SGX and weight was determined. **(D)** Systemic Leptin levels were quantified by ELISA after fasting in control and TDCA/valine-treated DIO and lean mice. **(E)** 2 g/kg glucose was injected following 8 hrs of daytime fasting. Blood glucose levels were assessed in blood samples utilizing a blood-glucose-meter. Results are representative of at least three independent experiments. Column plots display mean with standard deviation. Statistical significance was determined by using One-way-ANOVA. Asterisks indicate p-values * = p<0.05, **= p<0.01 and *** = p<0.001. Only significant values are shown (n=5-7 animals/group).

To further characterize the impact of TDCA/valine, we assessed subcutaneous and epididymal adipose tissue stores and found a significant decrease of both fat stores in TDCA/valine treated DIO mice comparable to effects observed in DIO mice that underwent SGx **(Fig. 3C)**.

In addition, we assessed whether treatment with TDCA/valine affected systemic levels of leptin as a main neuroendocrine peptide released by adipocytes(*24*). Indeed, our results indicated reduced systemic leptin levels in both, SGx and TDCA/valine-treated animals **(Fig. 3D)**.

It is well established that obesity promotes insulin resistance and type-2 diabetes(*2, 25*). Thus, to further explore the effects of TDCA/valine treatment on obesity-associated insulin resistance and T2D, we assessed the capacity for glucose tolerance. DIO mice treated with TDCA/valine displayed a complete reversal of obesity-related insulin resistance comparable to lean controls. Notably, beneficial metabolic effects subsequent to TDCA/valine were analogous to those observed in DIO mice following SGx (**Fig. 3E)**.

### Treatment with TDCA/valine induces weight loss through altered feeding behavior in the absence of physical dysfunction

Growing evidence suggests a decreased food intake as critical for the long-term weight reduction subsequent to SGx(*23*). Hypothesizing that TDCA/valine treatment may mimic SGx-induced weight loss, we next assessed calorimetric and metabolic parameters of DIO mice receiving daily injections of either TDCA/valine or PBS using a Comprehensive Laboratory Monitoring System (CLAMS). While cumulative energy intake and the hourly food intake significantly declined in TDCA/valine-treated animals compared to PBS-treated controls (**Fig. 4A, B**), hourly measured energy expenditure and the measured total activity did not change during treatment (**Fig. 4B, C**). Moreover, the respiratory exchange ratio and energy balance of TDCA/valine-treated obese animals significantly declined, supporting the significance of fatty acids for energy metabolism (**Fig. 4E, F**). Of note, read-outs had not been impacted by the circadian cycle as comparable results were obtained during light and dark periods. Intriguingly, our data on weight loss induced by TDCA/valine were neither linked to a typical food-seeking behavior in response to fasting, nor to unspecific toxic side-effects of the applied metabolites. Instead, treatment with TDCA/valine specifically targeted feeding behavior while leaving locomotor activity and energy expenditure unaffected thus generating a negative energy balance.

**Figure 4.**
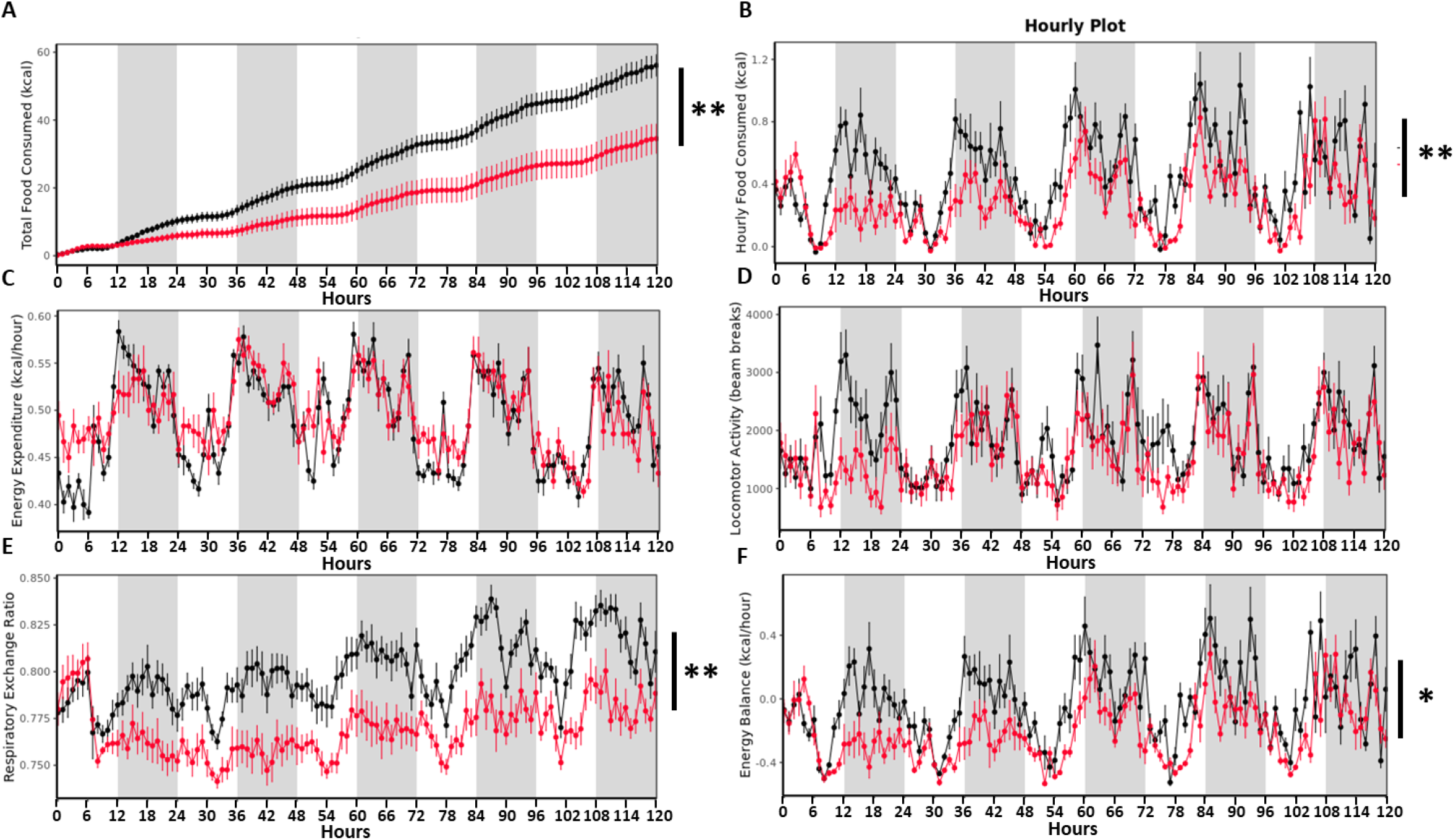
TDCA/valine-treatment induces weight loss through altered feeding behavior in the absence of reduced energy expenditure. 12 DIO mice (control=6, treatment=6) were placed into the Columbus Instruments Comprehensive Lab Animal Monitoring System (CLAMS) for 6 days. Time graphs represent hourly averages throughout the experiment. Shaded regions represent the 12-hr dark photoperiod. After one day of acclimation (not shown), injections of TDCA/valine were performed at 2pm for 5 days. This experiment monitored **(A)** cumulative energy intake **(B)** hourly food intake, **(C)** energy expenditure, **(D)** locomotor activity **(E)** respiratory exchange ratio, and **(F)** energy balance (energy intake minus energy expenditure). Results are representative of at least three independent experiments. Statistical significance was determined by ANOVA using total mass as the covariate. Error bars represent SEM. Asterisks indicate p-values *= p<0.05, **= p<0.01. Only significant values are shown. (n=6 animals/group).

### TDCA/valine-induced weight loss is mediated through suppression of hypothalamic levels of orexigenic MCH

After detecting reduced feeding behavior and preserved energy expenditure linked to TDCA/valine-associated weight loss, we next analyzed hypothalamic peptides that are involved in regulation of appetite and energy homeostasis while treating animals with TDCA/valine. To this end, we isolated hypothalamus tissue from acute brain slices and analyzed mRNA levels of neuroendocrine regulators (AgRP, CART, MCH, NPY, POMC) by RT-PCR, comparing expression in fed versus fasted animals. Our analysis revealed a substantial attenuation in the increase of melanin-concentrating hormone (MCH) with fasting following TDCA/valine treatment (**Fig. 5A**). Obese animals treated with PBS and fasted for 12 hours prior to procurement of hypothalamus tissue showed a pronounced increase in MCH levels, findings that were in line with the orexigenic effects of this neuropeptide. In TDCA/valine-treated obese animals, however, MCH-increase was not observed, suggesting an effect of TDCA/valine on the regulation of hypothalamic MCH expression **(Fig. 5A)**. Notably, MCH has been shown to centrally promote food intake, augment anabolic energy regulation(*26-28*) and increase body weight in experimental models(*29*).

**Figure 5.**
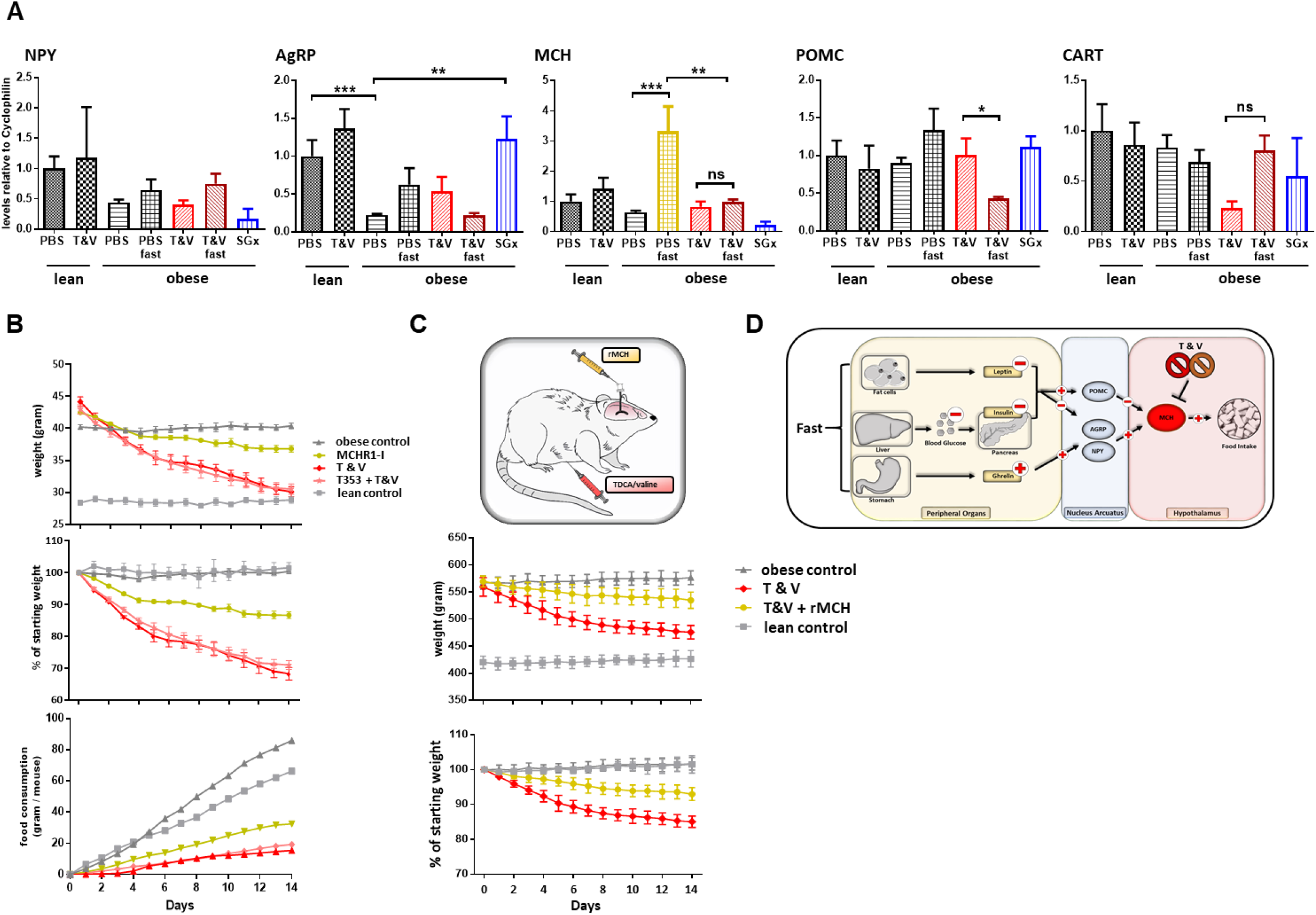
TDCA/Valine treatment acts through suppression of hypothalamic MCH levels (A) Lean and DIO mice were treated daily with either PBS or T&V. 2 Groups of T&V-treated DIO mice were subjected to 12h fasting before tissue procurement. After 2 weeks, all mice were sacrificed, hypothalamus tissue dissected and RNA levels of POMC, CART, NPY, AgRP and MCH measured by rtPCR. **(B)** DIO mice were subjected to daily *i*.*p*. injection of T&V, oral administration of MCHR1-I, or a combined treatment of both, T&V + MCHR1-I for a course of 2 weeks. Body weight and food consumption was measured and expressed as a contingency plot displaying total weight, percentage of starting weight and food consumption per mouse. **(C)** DIO rats were subjected to combined *i*.*p*. TDCA/valine injection and intracerebral administration of recombinant MCH and weight loss was monitored for 2 weeks. Results are representative of at least three independent experiments. Column plots display mean with standard deviation. Statistical significance was determined using One-Way-ANOVA. Asterisks indicate p-values *= p<0.05, **= p<0.01 and *** = p<0.001. Only significant values are shown. (n=5-7 animals/group).

To analyze the effect of TDCA/valine on MCH-regulated appetite and energy homeostasis in-vivo, we made use of a MCH receptor 1 inhibitor (MCHR1-I)(*30*). Treatment with MCHR1-I in DIO obese mice led to decreased food intake and subsequent weight loss (**Fig. 5B**). More importantly, simultaneous treatment with both MCHR1-I and TDCA/valine did not further reduce food intake or promote additional weight loss in DIO mice, suggesting that effects of TDCA/valine are, at least in part, mediated through suppression of hypothalamic levels of MCH.

Central administration of MCH has been reported to promote food intake in rats(*29*) while peripherally administered MCH has been shown to be unable to pass the blood-brain barrier(*31*). We thus used DIO Wistar rats that underwent cranial surgery, mounting a canula into the lateral cerebral ventricle, thus providing access for intracranial injections. To confirm a specific TDCA/valine-induced inhibition of MCH, we used a combinatorial treatment regimen of intraperitoneal administered TDCA/valine and centrally administered recombinant MCH. Consistent with our findings in mice, obese rats rapidly lost weight under TDCA/valine treatment within two weeks **(Fig. 5C)**. Strikingly, this effect was significantly diminished when co-administering recombinant MCH and TDCA/valine **(Fig. 5C)**, suggesting an inhibitory effect of TDCA/valine on MCH regulation **(Fig. 6)**.

**Figure 6.**
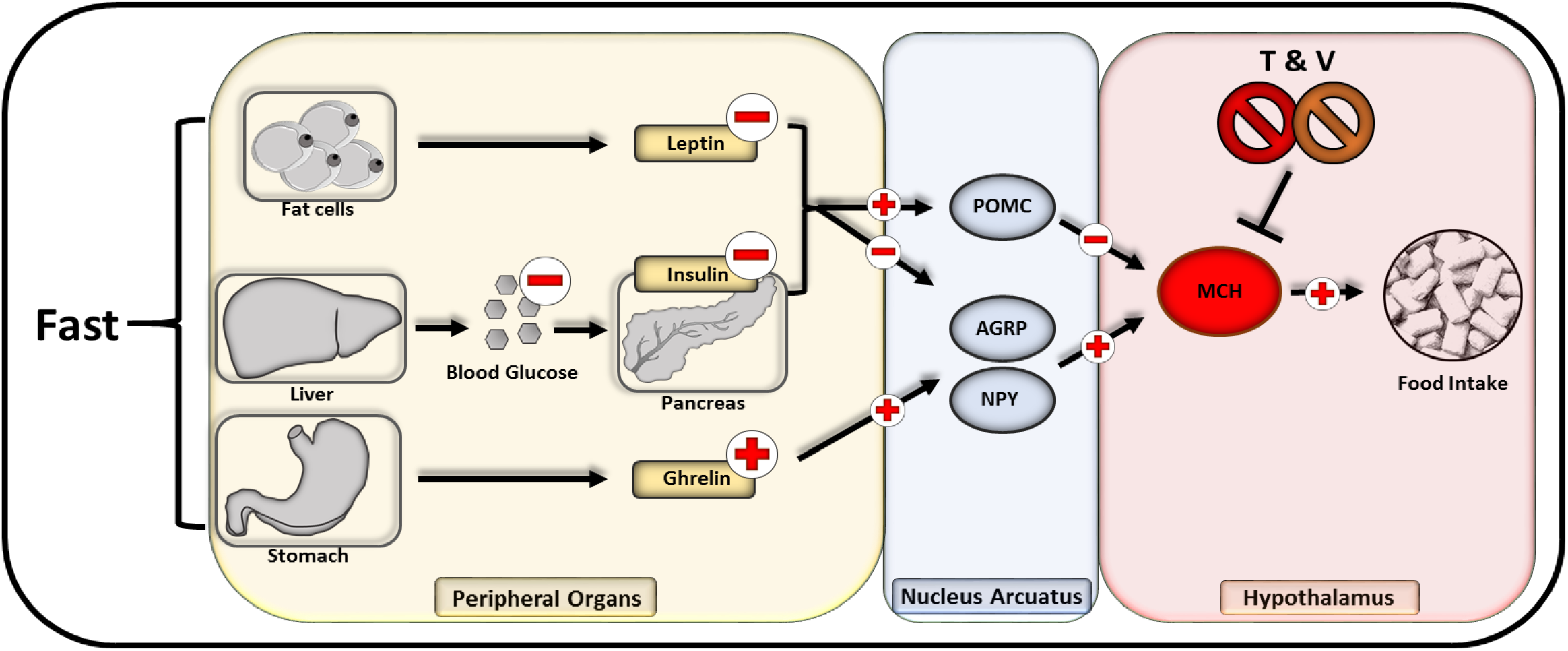
Flowchart of neuropeptide-mediated appetite regulation and TDCA/valine interaction.

## Discussion

Obesity has been associated with major metabolic changes that contribute to obesity-related disorders(*25*). In morbid obesity, bariatric surgery is often the only alternative to achieve sustained weight loss. More importantly, among various bariatric procedures, SGx has been shown to induce favorable metabolic changes(*18, 32, 33*) while improving insulin sensitivity and glycemic control. Although SGx has made significant advances in reversing obesity, many risks are associated with this surgical procedure including excessive bleeding, infection, blood clots and leaks resulting from surgery, thus urging for alternative, non-invasive approaches.

Consistent with previous reports, we observed a prominent and durable weight loss following SGx. More importantly, we identified restored systemic levels of the bile acid, TDCA, and the branched-chain amino acid, L-valine in DIO mice that underwent SGx in contrast to DIO control mice in which those metabolites had been depleted. TDCA and L-valine levels demonstrated a similar pattern in obese patients that underwent SGx, confirming the clinical relevance of our findings.

Recently, bile acid signaling has been identified as a mechanistic underpinning of weight loss subsequent to SGx(*18*). Indeed, durable weight loss and improved glucose tolerance are mediated through bile acid-induced FXR signaling, affecting overall food intake and feeding behavior(*18*). Thus, weight loss subsequent to SGx may only in part be communicated through the restricted gastric capacity. Moreover, re-feeding mice that underwent SGx with a HFD revealed that these animals gained weight without reaching pre-surgery obesity, supporting a relevant role of appetite behavior in addition to a modified metabolism(*34*). In contrast, diet-restricted DIO mice, restored lost body weight by exhibiting hyperphagia, fully regaining their bodyweight with the end of the diet(*35*). Hence, a decreased food intake is likely to constitute a primary impetus for the long-term weight loss observed after SGx (*23*). In support, pair-feeding studies have shown that rats restricted to the same daily caloric intake as SGx animals exhibited comparable weight loss(*23*).

These findings are in line with our study, suggesting that durable weight loss and amelioration of obesity associated disorders after SGx are mediated through both, molecular and central appetite-regulating effects of TDCA/valine. Indeed, our results indicated, that TDCA/valine-treated animals had a markedly reduced food intake without altered energy expenditure or locomotor activity, thus excluding toxic side effects of TDCA/valine as a potential confounder of appetite reduction. Notably, TDCA/valine administered to lean animals did not lead to weight loss, emphasizing on appetite regulating effects only in obese animals while excluding inflammatory processes as confounding side effects. Therefore, our observation that TDCA/valine treatment mimicked metabolic changes that result from SGx are supported by our finding that decreased food intake is a cardinal feature of SGx-induced weight loss.

Within the last decades increased levels of branched-chain amino acids (BCAA) levels have been associated with T2D and obesity derived complications(*36*). Notably, various studies have suggested increased BCAA levels to be a consequence of obesity rather than its cause. Indeed, mitochondrial impairment and obesity derived low grade inflammation has been associated with decreased levels of enzymes that degrade BCAA (*37, 38*). Moreover, a recent study employing a combination of thiazolidinediones and metformin failed to demonstrate decreasing systemic BCAA levels despite an increased insulin sensitivity(*39*). We thus speculate that the weight reducing effect of L-valine in combination with TDCA is not impacting obesity directly but rather exerts effects through a reduction of appetite.

Supporting our results, BCAA have shown beneficial effects on weight reduction. Supplementation of leucine to drinking water of DIO mice reduced body weight by 32% over 10 weeks(*40*). Furthermore, leucine administration has been shown to induce a partial resistance to diet-induced obesity that resulted into reduced hyperglycemia and hypercholesterolemia in DIO mice(*40*). The activation of Pro-opiomelanocortin (POMC) neurons in the nucleus arcuatus of the mediobasal hypothalamus(*41*) alters the feeding behavior of DIO mice and has been proposed as a critical mechanistic effect. Activation of mammalian target of rapamycin (mTOR), a signaling pathway integrating sensing of the energy status and growth and proliferation, for instance has been identified as a mechanistic underpinning of BCAA derived weight loss. Notably, intracerebroventricular administration of leucine decreased food intake leading to body weight loss in DIO rats through mTOR activation of POMC neurons in the nucleus arcuatus while rapamycin, a potent mTOR inhibitor abolished those effects(*42*). In the current study, we could not detect significant differences in the expression of POMC in obese and TDCA/valine treated animals that had fasted prior to hypothalamic procurement. However, POMC-neurons exhibit inhibitory effects on neurons of the lateral hypothalamic area in which the orexigenic peptide MCH is localized(*43*). It is well-established that MCH regulates feeding behavior, energy balance and promotes food intake(*29*). In line with those observations, we detected a dramatically augmented MCH expression in obese animals upon fasting that has been absent in both fasted TDCA/valine-treated animals and animals that had been kept on HFD and subjected to TDCA/valine administration. Since MCH is an essential mediator of appetite regulation(*44*), we hypothesized a potent effect of TDCA/valine on MCH suppression. Consistently, the combinatorial administration of a selective MCHR1 antagonist(*45*) and TDCA/valine failed to elicit additional weight loss, suggesting that TDCA/valine acts at least in part through inhibiting MCH. To confirm this hypothesis, we next made use of DIO Wistar rats that had a canula placed into the lateral cerebral ventricle. As in our mouse model, our data indicated that rats subjected to HFD exhibited a dramatic weight loss when subjected to TDCA/valine. Moreover, upon central administration of recombinant MCH, the effects of TDCA/valine treatment were significantly diminished, suggesting a direct effect of TDCA/valine onto the appetite regulating MCH network. Of note, TDCA/valine-derived weight loss had not been fully reversed subsequent to intrathecal MCH administration. Thus, model-dependent restrictions in MCH application or dosing aspects may be of relevance. In support, other studies have reported a decreased intensity of MCH-promoted food intake over the course of 24 hours with a peak efficiency 2 hours after administration(*29, 46*). Moreover, by restoring the bile acid pool(*18*) through TDCA administration, FXR-signaling is likely to constitute an additional pathway mediating metabolically derived weight loss. While our studies show that the weight reducing effects of TDCA/valine are communicated, at least in part, through MCH, additional pharmacological and mechanistic studies of TDCA/valine-mediated impact on hypothalamic neuropeptide expression will be necessary. Moreover, future studies will need to determine whether TDCA/valine administration may exert beneficial effects through elevated serum or cerebrospinal-fluid (CSF) levels and if additional metabolic effects altering mitochondrial metabolism and potentially affecting other BCAA levels are in effect.

In summary, our results add critical novel data on mechanisms and consequences of surgery induced weight loss. More importantly, and of translational relevance, we introduce TDCA and L-valine as novel agents for weight loss and appetite regulation.

## Materials and Methods

### Animals

Animal use and care were in accordance with institutional and National Institutes of Health guidelines. Diet induced obese (DIO) C57BL/6 mice and lean littermates were purchased from Taconic (Taconic Farms Inc. Germantown, NY) for all studies. The study protocol was approved by the Brigham and Women’s Hospital Institutional Animal Care and use Committee (IACUC) animal protocol (animal protocol 2016N000371). Obesity was induced by feeding animals ad libitum with a high fat diet (HFD) that provides 60% of total energy as fat (D12492 diet, Research Diets Inc. New Brunswick, NJ) starting at 6 wks of age for a duration of 12 wks. Wistar rats used for intracerebral administration of recombinant MCH were group-housed and fed D12492 for 12 weeks (60% k/cal diet). Rats subsequently underwent cranial surgery placing a resealable canula in the lateral cerebral ventricle. All animals were maintained in specific pathogen-free (SPF) conditions at the Brigham and Women’s Hospital animal facility in accordance with federal, state, and institutional guidelines. Animals were maintained on 12-h light, 12-h dark cycle in facilities with an ambient temperature of 19–22 °C and 40–60% humidity and were allowed free access to water and standard chow. Euthanasia was performed by cervical dislocation following anesthesia with isoflurane (Patterson Veterinary, Devens, MA, USA).

### Bariatric surgery

A gastric sleeve was created along the lesser curvature by transecting the stomach. The sleeve was then hand sewn, using an 8-0 continuous Prolene suture. Sham animals underwent a laparotomy, the stomach was isolated and blunt pressure was applied with forceps for corresponding durations.

### Metabolic experiments

- Conservative weight loss DIO-C57BL/6 mice obese animals that were on a HFD (12492, Research Diets INC.) at 6 wks of age for a duration of 12 wks were then switched to a normal chow diet (NCD) and consecutive body weight was assessed.
- Intraperitoneal glucose tolerance test (IPGTT) Glucose (2 g/kg, Sigma Aldrich, St. Louis, MO) was injected for glucose tolerance testing which was conducted by 8 hrs of daytime fasting. Blood glucose was monitored using an Accu-Check Aviva Plus blood glucose meter (Roche, Plainfield, IN).
- Metabolomic treatment Taurodesoxycholic acid (TDCA) (50mg/kg) and L-Valine (200mg/kg) (both from Sigma Aldrich, St. Louis, MO) were dissolved in sterile ddH_2_O and simultaneously administered by intraperitoneal injection.
- MCHR1-I administration MCHR1-I was obtained from Takeda, Chuo-ku, Tokyo, Japan. MCHR1-I was administered at 10mg/kg via oral gavage. As vehicle solution 0.5% Methocel, 0.1% Tween 80, 99.4% distilled water was used. For combinatory treatment of MCHR1-I and TDCA/valine, mice were additionally injected intraperitoneally with TDCA/valine.
- rMCH administration Recombinant MCH (Cayman Chemical) was dissolved in sterile ddH_2_O (1ug/ul) and 5μl was administered by intracranial injection over 30 seconds into the lateral ventricle.

### Indirect Calorimetry

12 DIO mice (control=6, treatment=6) were placed into the Columbus Instrument Comprehensive Lab Animal Monitoring System and maintained for 6 days. They were kept on HFD at 22C +/-1°C ambient temperature for the duration of the experiment. After one day of acclimation, injections of TDCA/valine were performed at 2pm each day for 5 days. Time graphs represent hourly averages throughout the experiment. Bar graphs correspond to the total, light, and dark cycles (12-hour cycles beginning and ending at 6am and 6pm). Error bars represent SEM. Student t-tests were performed on all bar graphs.

### Human samples

Serum samples from patients prior to and 3 months post sleeve gastrectomy were obtained with approval of the Brigham and Women’s Hospital (BWH) Institutional Review Board and through cooperation with Dr. Eric G. Sheu and the Center for Metabolic and Bariatric Surgery at BWH. Informed consent was obtained from all patients and samples were collected following BWH ethical regulations. Whole blood samples were obtained at routinely scheduled pre-operative and post-operative appointments, centrifuged to obtain serum and then stored at −80°C up until metabolite measurements and data analysis exactly as described above for murine samples.

### Metabolite measurements by LC–MS/MS and data analysis

Whole blood samples were centrifuged at 13.000g for 10 min at 4°C, and 200µl of the supernatant were saved; 800 µl of cooled methanol (−80°C) were added to the supernatant for a final 80% (vol/vol) methanol solution. Samples were incubated for 6 hours at −80°C, and then centrifuged at 13,000×g for 10 min at 4 °C. Supernatants were collected, dried in a SpeedVac (Savant AS160, Farmingdale, NY), and stored at −80°C until analysis. Each sample was resuspended in 20 μl of LC/MS grade water and then analyzed with a 5500 QTRAP, a hybrid triple quadrupole/linear ion trap mass spectrometer, using a quantitative polar metabolomics profiling platform with selected reaction monitoring (SRM) that covers all major metabolic pathways. Quantitative analysis of 260 detected metabolites was performed utilizing the web-based MetaboAnalyst 3.0 software.

### Dissection of hypothalamus tissue, RNA extraction, realtime PCR

Mice were assigned to the following experimental groups, with n=6 in each group: lean PBS-treated mice, lean TDCA/valine treated mice, DIO-obese PBS treated mice, DIO-obese PBS treated mice fasted for 12h before tissue procurement, DIO-obese TDCA/valine treated mice, DIO-obese TDCA/valine treated mice fasted for 12h before tissue procurement, DIO obese mice undergoing sleeve gastrectomy. Cage beddings were pooled and redistributed at days −6, −4, and − 2 to normalize microbial flora among experimental groups. At day 0, treatment began and sleeve gastrectomies were performed. Mice were treated for 13 days, with procurement of tissues performed on day 14. Animals were anaesthetized with ketamine and sacrificed by decapitation. Brains were removed, and hypothalamus tissue was dissected and flash-frozen in liquid nitrogen. RNA was isolated using Direct-zol RNA MiniPrep kit (Zymo Research, Irvine, CA). cDNA was made from isolated RNA using oligo (dt), random hexamer primers and reverse transcriptase QuantiTech RT Kit (Qiagen, Germantown, MD). Quantitative PCR was performed using the 7800HT (Applied Biosystems, Foster City, CA) thermal cycler and SYBR Green master mix (Applied Biosystems). Relative mRNA abundance was calculated and normalized to levels of the housekeeping gene cyclophilin.

### Statistics

Unless otherwise specified in figure legends, comparisons between experimental groups were performed using Student’s t test. Survival curves were compared by using the log rank test. When applicable, mice were randomly assigned to treatment or control groups. All results were generated using GraphPad Prism software (San Diego, CA). A p-value of 0.05 was considered statistically significant.

## Acknowledgements

The authors wish to express their appreciation and gratitude to Dr. Maratos-Flier, Division of Endocrinology, Diabetes and Metabolism, Beth Israel Deaconess Medical Center, Boston, MA, USA and Novartis Institute for BioMedical Research, Cambridge, MA, USA for expert advice and helpful discussions.

## Author contributions

M.Q., J.I., T.H. performed experiments, analyzed data and wrote the manuscript. H.R.C.B., Y.N., F.K. performed experiments. H.U., R.M. supported experiments. A.E. and S.G.T. designed experiments and wrote the manuscript.

